# Distinct Roles of Working Memory and Inhibitory Control in Speech-in-Noise Perception

**DOI:** 10.64898/2026.09.17.752459

**Authors:** Sharadhi Bharadwaj, Catherine Dymowski, Erin Batik, Grace Caplan, Courtney Shepler, Amanda Yagan, Xingyu Zhang, Hari Bharadwaj, Aravindakshan Parthasarathy

**Author notes:** **Corresponding author**: Aravindakshan Parthasarathy, Department of Communication Science and Disorders University of Pittsburgh, Pittsburgh, PA 15213.

## Abstract

Difficulty perceiving speech in noise is the primary complaint of patients seeking hearing healthcare, even absent overt losses in audiometric thresholds. Yet current clinical practice largely focuses on sensory function while ignoring the potentially outsized effects of executive function on everyday suprathreshold listening. Speech perception in noise depends on the dynamic interplay between the fidelity of sensory coding along the auditory pathway and higher-order top-down cognitive processes that support goal-directed listening, both of which can independently change across the lifespan. Here we examined how two subcomponents of executive function, working memory and inhibitory control, vary across the adult lifespan and contribute to speech-in-noise performance. Two hundred participants aged 18 to 73 with normal hearing thresholds were tested on a validated online auditory psychophysics platform. Working memory was assessed with an n-back task and inhibitory control with an auditory Stroop task. Participants then performed two speech-in-noise tasks: a modified rhyme test emphasizing sensory coding through energetic masking at the cochlea, and a matrix sentence task emphasizing executive function through informational masking. Age significantly affected performance on both tasks. Aging was associated with reduced inhibitory control (increased Stroop cost) but no significant effect on working memory. Regression analyses revealed that both working memory and inhibitory control predicted performance on the informational masking task, whereas only inhibitory control predicted performance on the energetic masking task. These findings underscore the critical role of executive function in everyday suprathreshold listening and support including cognitive measures in clinical assessments of hearing difficulties.

## Introduction

Difficulty understanding speech in noise is among the most common complaints of adults seeking hearing healthcare, even when audiometric thresholds fall within the clinically normal range (Cancel et al., 2023; Hind et al., 2011; Parthasarathy et al., 2020; Spehar & Lichtenhan, 2018). Yet, current clinical practice remains focused largely on sensory function and audibility. This approach inadequately captures the experiences of these listeners, whose challenges often lie beyond audibility *per se*.

Additional sources of these suprathreshold deficits remain an active area of research. Possible peripheral sensory and neural mechanisms described so far include cochlear synaptopathy or a loss of synaptic connections between inner hair cells and the auditory nerve fibers that transmit the sound signal to the brain (Kujawa & Liberman, 2009), distorted tonotopy or a change in tonotopic tuning properties in the auditory nerve (Henry et al., 2016; Parida & Heinz, 2022), and altered neural synchrony at the level of the auditory nerve (Dias et al., 2024; Xing et al., 2012). However, successful speech perception in noise relies not only on the fidelity of sensory encoding but also on suprasensory, “top-down” cognitive processes that support goal-directed behavior(Kuchinsky et al., 2013; McHaney et al., 2024; Parthasarathy et al., 2020; Peelle, 2018; Winn et al., 2015; Zink et al., 2024).

These top-down factors facilitate the listener’s ability to allocate attention, maintain relevant information, and suppress distractors. These processes, collectively referred to as *executive functions*, are increasingly recognized as key contributors to real-world listening.

Two subcomponents of executive functioning, working memory, and inhibitory control are especially relevant to speech understanding in noise. Working memory supports the temporary maintenance and manipulation of auditory information, allowing listeners to integrate degraded acoustic input over time to derive meaning (Akeroyd, 2008; Rönnberg et al., 2013). Age related declines in working memory have been linked to poorer speech in noise performance (Meister et al., 2013). Furthermore, studies in individuals with hearing impairment and hearing-aid users show that higher working-memory scores predict better speech-in-noise outcomes (Ng et al., 2013; Rudner et al., 2011). Inhibitory control supports suppression of competing distractors or auditory information, enabling listeners to maintain focus on task-relevant speech information (Nagaraj, 2021). Behavioral and neurophysiological evidence indicates that age-related declines in inhibitory control contribute to poorer speech-in-noise perception, independent of peripheral hearing sensitivity (Brungart, 2001; Gómez-Álvarez et al., 2023; Hasher & Zacks, 1988). Such deficits may reflect both cognitive inefficiency and age-related reductions in neural inhibition along the auditory pathway, potentially compounding sensory degradations such as cochlear synaptopathy (Gómez-Álvarez et al., 2023).

To disentangle the relative contributions of executive and sensory mechanisms, speech-in-noise performance can be examined under masking conditions that differentially tax auditory versus cognitive processes. Energetic masking (EM) occurs when noise overlaps both spectrally and temporally with the target, interfering at the auditory periphery. Therefore, EM is thought to primarily reflect the sensory encoding issues (Shinn-Cunningham, 2008). In contrast, Informational masking (IM) occurs when maskers are non-overlapping in time or frequency but rather contain meaningful information or speech-like content that interferes at the central auditory and/or higher cognitive levels. Successful performance in IM tasks require the listener to suppress irrelevant speech and attend to the target speech stream, thereby placing demands on both inhibitory control and working memory.

Here, we investigate the contributions of working memory, inhibitory control, and age to speech perception in noise under both energetic masking (EM) and information masking (IM) tasks. A large online sample of adults spanning a broad age range completed auditory versions of the *n*-back task and the Stroop task to assess working memory and inhibitory control, respectively. Speech-in-noise performance was measured using two complementary paradigms: a Modified Rhyme Test (MRT) with inharmonic tone maskers to probe performance under energetic masking (House et al., 1963; Stone & Moore, 2014), and a Matrix Sentence Task for probing performance under informational masking (Kidd et al., 2008; Swaminathan et al., 2015). We hypothesized that (1) age would significantly predict performance in all tasks, and (2) measures of executive function would account for significant variance in IM task performance, even after controlling for age but not in the EM task performance.

## Methods

### Participants

Data were collected from 200 participants (18-73 years old; mean age= 39±12; 95 female, 92 male, 2 other) recruited anonymously through Prolific, an online research platform. All participants were residents of US or Canada and were native speakers of North American English. Participants reported normal hearing (confirmed through an online speech-based screener described below), normal or corrected to normal vision, and no neurological disorders.

Participants were instructed to use stereo headphones in a quiet room. A headphone screening procedure confirmed that they could detect the auditory stimuli at comfortable levels. All procedures were approved by the institutional review board of the University of Pittsburgh (STUDY24050210).

### Online hearing screener

Prior to the main experiment, all participants completed a two-step screening procedure to ensure they were using headphones and had putative normal hearing, following the framework established by Mok et al. (Mok et al., 2024a). First, the participants completed a headphone screening task, adapted from Woods et al. (Woods et al., 2015) where the participants were presented with a series of tones and required participants to identify the softest tone. One of these tones served as an ‘anti-phase decoy,’ presented with the left and right channels 180° out of phase. When played over loudspeakers, the out-of-phase signals from the left and right channels physically mix in the air, resulting in destructive interference that significantly reduces the perceived loudness of this tone, often making it sound like the softest tone. In contrast, headphones physically isolate the left and right channels, preventing this acoustic cancellation. Therefore, only participants wearing headphones perceive the decoy at its full, high intensity and correctly identify the intended target (−6 dB) as the softest tone. Once participants completed the headphone check, they were prompted to complete a hearing screening task. The participants completed a modified rhyme test (MRT). The target word was presented with a carrier phrase “Please select the word______” in a four-talker babble at different SNRs. For each trial, participants were required to identify the target from a set of six rhyming alternatives (6-alternative forced-choice). To be able to participate in the main experiment, participants were required to meet specific performance criteria derived from meta-analytic data on normal-hearing listeners: 100% accuracy at +10 dB SNR, ≥83% accuracy at +5 dB SNR, and ≥75% accuracy at 0 dB SNR (Mok et al., 2024b).

### Speech-in-noise tasks

Speech perception in noise was assessed using two paradigms: a Modified Rhyme Test (MRT), which predominantly engages sensory coding under *energetic masking* conditions, and a Matrix Sentence Task, which stresses cognitive control under *informational masking* conditions.

Modified Rhyme Test (MRT): In the MRT, each target word was masked by an inharmonic tone complex and preceded by the carrier phrase *“Please select the word*.*”* Participants selected the correct word from six visually presented rhyming monosyllabic options (e.g., *went, sent, bent, dent, tent, rent*; Fig. 1C) (House et al., 1963; Stone & Moore, 2014). Stimuli were presented at six fixed signal-to-noise ratios (SNRs): −24, −18, −12, −6, 0, and +6 dB, with 12 trials per SNR. The masker level was held constant while the target speech level was varied to achieve the desired SNR.

**Figure 1.**
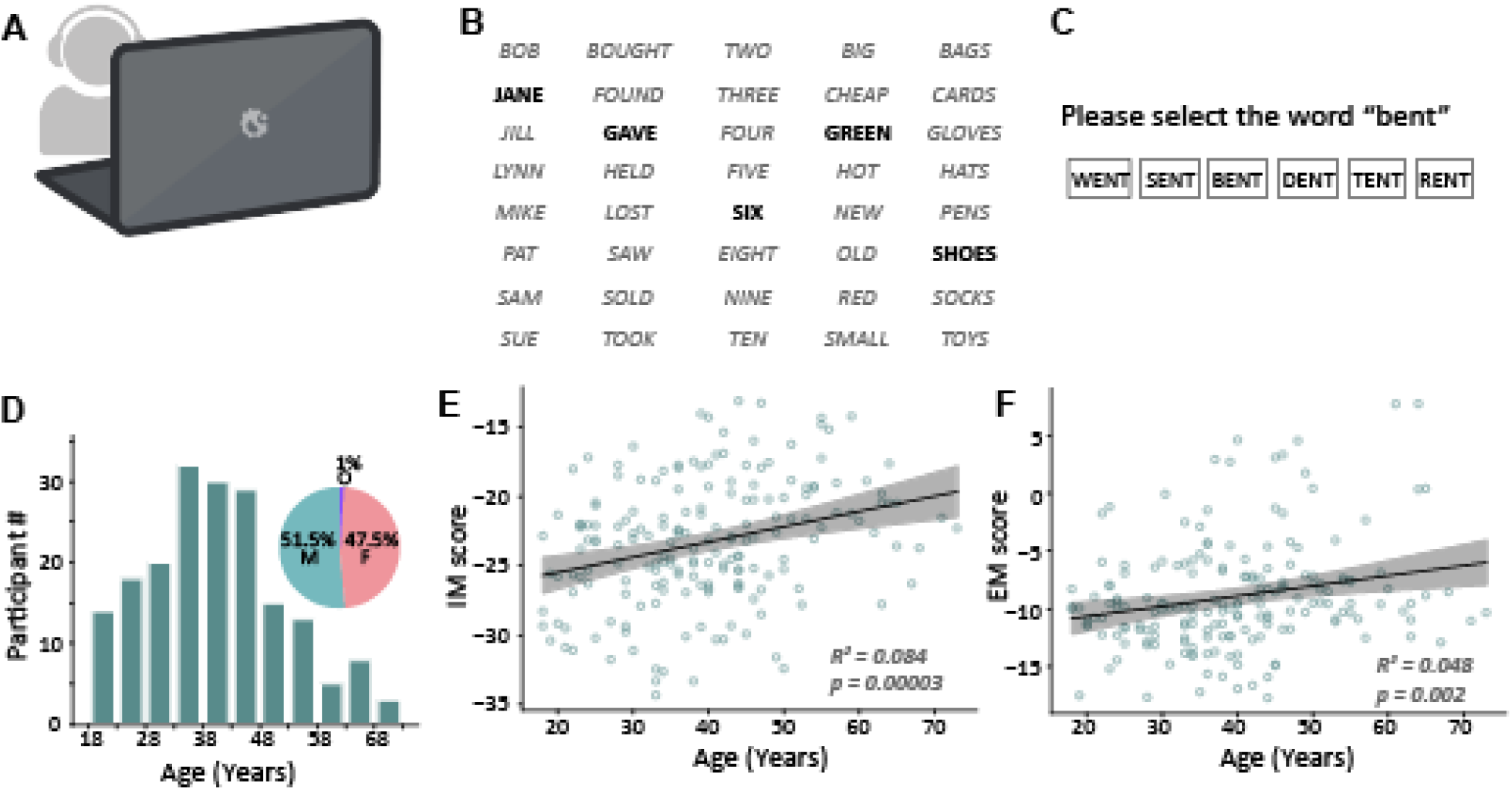
Experimental paradigm and age-related decline in speech-in-noise perception. (A) Schematic of the web-based remote testing setup; participants completed tasks at home using personal computers and headphones. (B) Schematic of the Matrix sentence task used to assess Informational Masking (IM). Participants identified a target sentence (starting with the call sign “Jane”) presented amidst two competing female talkers. (C) Schematic of the Modified Rhyme Test (MRT) used to assess Energetic Masking (EM). Participants identified a target word masked by an inharmonic tone complex from six rhyming alternatives. (D) Histograms showing the distribution of age, and pie chart for sex of the study cohort(inset) (N=200). (E) Performance on the IM task as a function of age. The solid line represents the linear regression fit, and the shaded region represents the 95% confidence interval. IM scores significantly declined with age (p<0.001). (F) Performance on the EM task as a function of age, showing a significant decline in performance with increasing age (p=0.002)

Matrix sentence task – The Matrix Sentence Task consisted of syntactically correct but semantically unpredictable sentences, each comprising five words drawn randomly from five categories - name, verb, number, adjective, and noun (8 options per category; Fig. 1B), for a total of 40 unique words (Kidd et al., 2008). Stimuli were produced by mixing three spatially separated female talkers, with one designated as the target. The target sentence always began with the name *“Jane”* and occurred diotically. The two other talkers produced masker sentences composed of different word combinations and presented from virtual azimuthal angles of +15 degrees and –15 degrees using non-individualized head-related transfer functions (HRTFs). The stimuli were presented at SNRs of −35, −25, −15, −5, 0, and +5 dB, with 10 trials per SNR. The subjects were scored out of 40 (4 words per trial) for each SNR. As with the MRT, masker level was fixed and target level varied to manipulate SNR.

### Executive functioning measures

Two auditory tasks were used to assess subcomponents of executive function: working memory (auditory *n*-back) and inhibitory control (auditory Stroop).

Auditory N-back task – The auditory N-back task was adapted from Kane et al 2007. (Kane et al., 2007) where participants heard sequences of spoken English letters and were instructed to press a designated key whenever the current letter matched the one presented *n* steps earlier in the sequence. The stimulus consisted of a subset of letters from English alphabet which were phonemically distinct - B, F, K, H, M, Q, R, X. Each participant completed 1-back, 2-back and 3-back conditions where 36 tokens were presented for each N-back condition of which each had 12 targets. The stimulus was spoken by a single female talker, and the intensity was set by the participants at a comfortable listening level. The interstimulus interval (ISI) between the spoken letters was 600ms. The participants responded by selecting if the token presented was a ‘target’ on the computer if the stimulus matched the one presented ‘N’ steps before or ‘non-target’ otherwise. Performance was quantified as *d′* (sensitivity index) for each *n*-level. Because performance was near ceiling for 1- and 2-back conditions, only 3-back *d′* values were included in subsequent analyses.

Auditory Stroop task – The auditory stroop task was adapted from Knight and Heinrich et al 2017 (Knight & Heinrich, 2017) where participants judged the pitch of a spoken word, while disregarding its semantic meaning. Words (“high” or “low”) were spoken by one of two male talkers at either a high (200–225 Hz) or low (75–95 Hz) fundamental frequency. Stimuli were either congruent (e.g., the word “high” spoken with high pitch) or incongruent (e.g., “high” spoken with low pitch) with 40 trials randomized in each condition. Participants responded as quickly and accurately as possible by pressing one of two keys corresponding to high or low pitch. Response times (RTs) were measured from stimulus offset for both congruent and incongruent trials.

### Data Analysis

For each participant, psychometric functions were fitted to performance across SNR levels for both speech-in-noise tasks. The SNR corresponding to 70% correct performance was estimated as the speech-in-noise threshold.

For the *n*-back task, sensitivity (*d′*) was computed separately for each *n*-level. Because 1- and 2-back tasks showed ceiling effects, only 3-back *d′* values were used in further analyses.

For the Stroop task, Stroop cost was calculated as the percent increase in mean RT for incongruent relative to congruent trials:

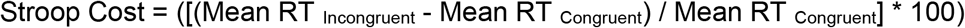

Linear regression analyses assessed relationships between age and performance on the speech-in-noise and executive-function tasks. Hierarchical multiple regression models were then used to test whether executive function measures explained additional variance in speech-in-noise performance after accounting for age. Age was entered in the first step, Stroop cost in the second, and 3-back d′ in the third. Separate models were run for energetic masking (MRT) and informational masking (Matrix Sentences) conditions.

All analyses were performed in Python using the *statsmodels* package (Seabold & Perktold, 2010).

## Results

### Older age is associated with poorer performance on both informational and energetic masking tasks

Regression analyses revealed significant positive associations between age and both informational masking (IM) and energetic masking (EM) thresholds (Fig. 1E-F). Age significantly predicted performance on the IM task (Fig 1E, p<0.001), though it only explained a small proportion of the variance (R^2^ = 0.084). Similarly, age was also a significant predictor for the EM task (R^2^ = 0.048, p = 0.002).

These findings indicate that normal hearing older adults exhibit modest but reliable declines in speech-in-noise performance across both masking conditions.

### Older age is associated with reduced inhibitory control but not working memory performance

Mean *d′* values in the auditory *n*-back task decreased systematically with increasing task difficulty: 9.29 for 1-back, 5.38 for 2-back and 1.45 for 3-back (Fig 2B). A linear regression examining the relationship between age and 3-back *d′* revealed no significant association (*p* = 0.07, *R*^*2*^ = 0.013; Fig. 2C), indicating that working-memory performance was relatively preserved across the adult lifespan.

**Figure 2.**
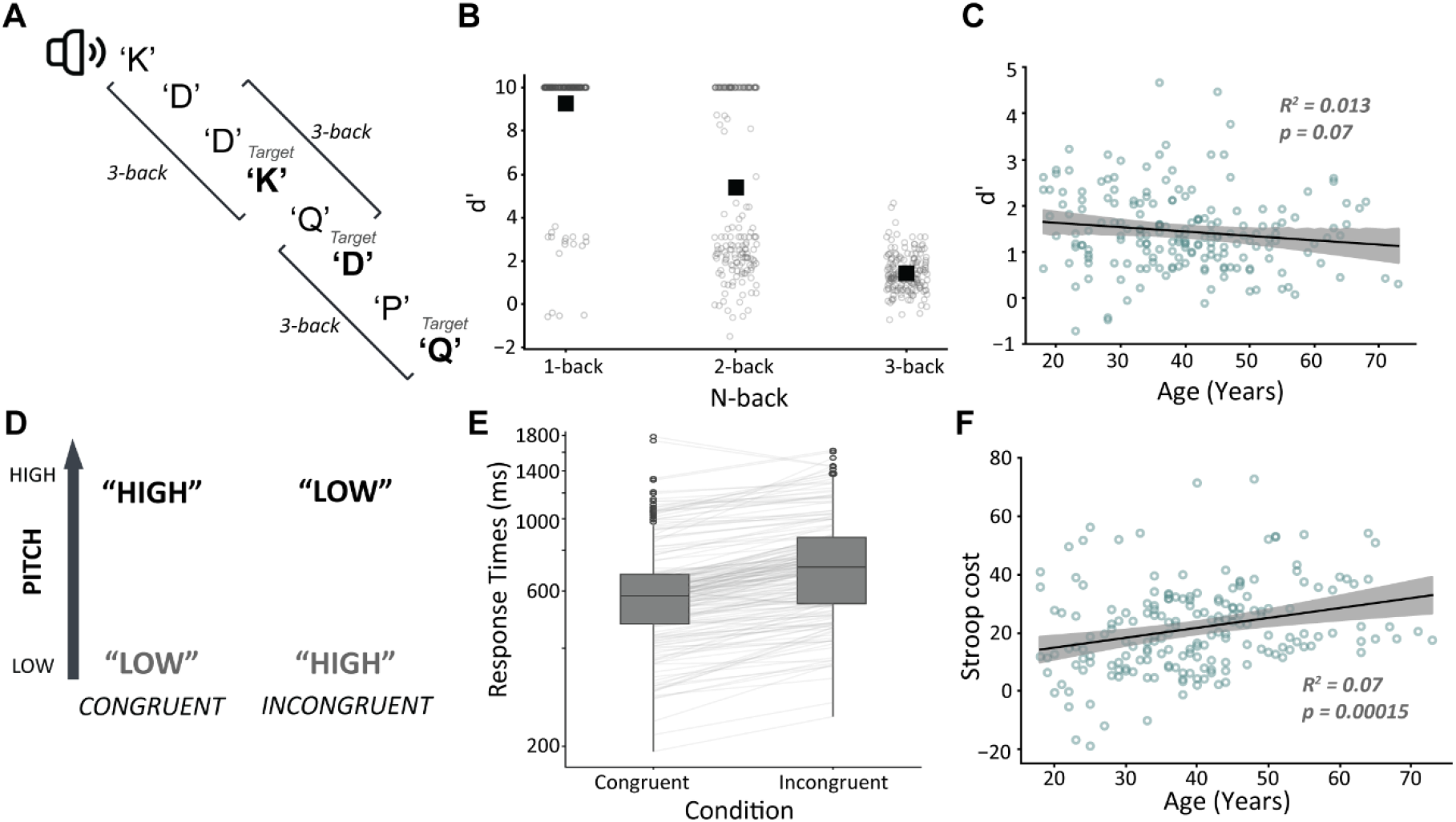
Age-related changes in executive function. – A) Schematic of the auditory N-back task used to assess working memory. Participants were required to detect a match between the current stimulus and the one presented N steps earlier. (B) Distribution of sensitivity (d’) scores for the 1-back, 2-back, and 3-back conditions (C) Performance on the 3-back task (d’) as a function of age. The regression analysis revealed no statistically significant decline in working memory performance with age (p=0.07). (D) Schematic of the auditory Stroop task used to assess inhibitory control. Participants judged the pitch of the spoken word while ignoring its semantic content. (E) Mean response times for congruent (pitch matches word) and incongruent (pitch mismatches word) trials across all participants, demonstrating the expected Stroop interference effect. (F) Inhibitory control performance as a function of age, showing a significant decline (increased Stroop cost/response time) with increasing age (p<0.001).

In contrast, performance on the auditory Stroop task showed clear age-related changes. Response times were longer for incongruent than for congruent trials (Fig. 2E), consistent with a Stroop interference effect. A linear regression revealed a significant positive relationship between age and Stroop cost, with age explaining approximately 7% of the variance (p < 0.001, R^2^ = 0.07; Fig. 2F).

These findings suggest that inhibitory control declines with age, whereas working-memory performance remains relatively stable across adulthood.

### Executive function measures selectively explain variance in informational masking and energetic masking performance

Hierarchical regression analyses were conducted to assess whether inhibitory control (Stroop cost) and working memory (n-back d′) accounted for additional variance in speech-in-noise performance after controlling for age (Fig. 3).

**Figure 3.**
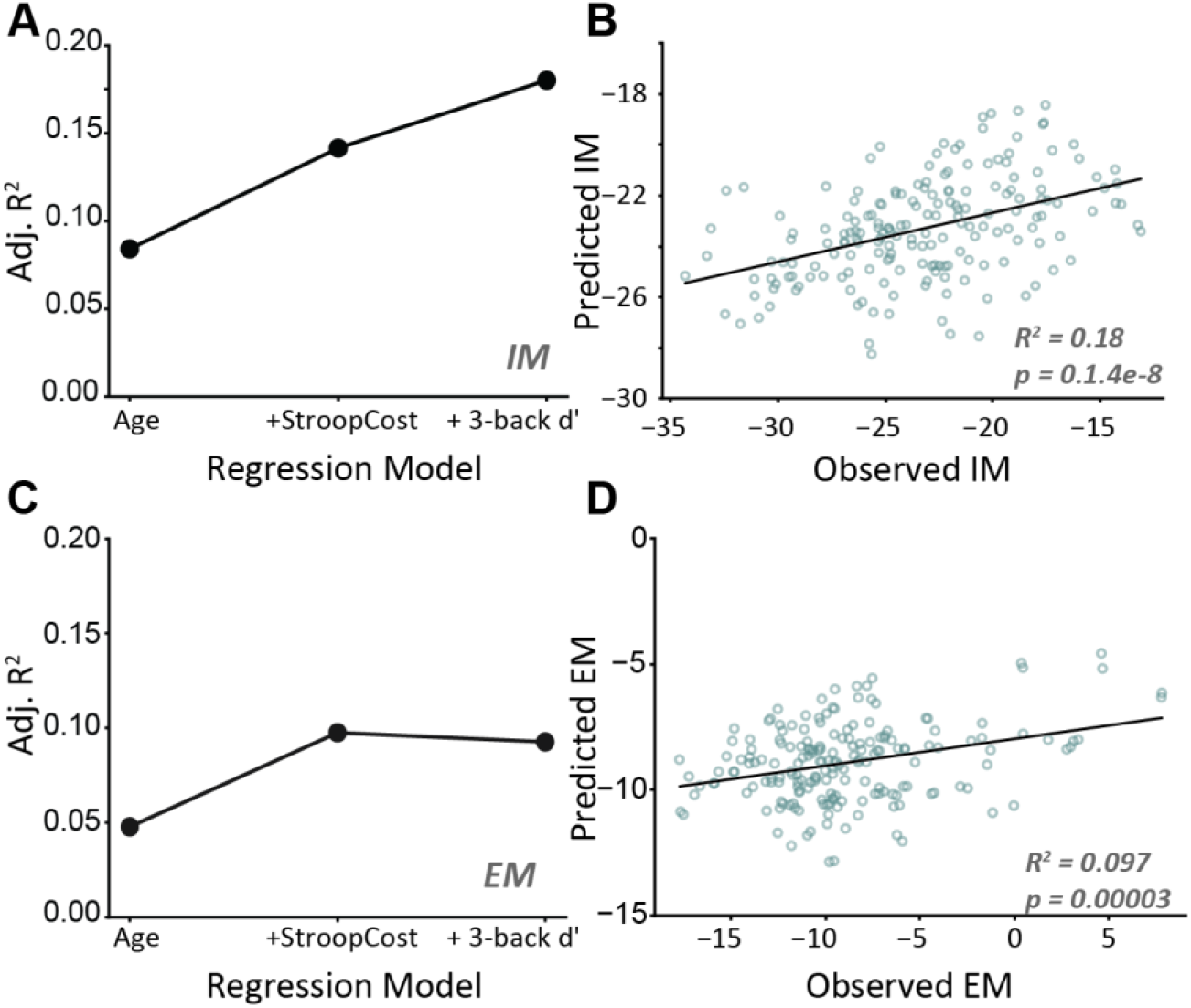
Hierarchical regression analysis of speech-in-noise performance. (A) The Adjusted R^2^ of the regressions models to predict IM performance. The variance explained improves with addition of each variable (age, age+StroopCost, age+StroopCost+3-back d’). (B) Predicted versus observed scores for the Informational Masking (IM) task. The final model, which included Age, Inhibitory Control (Stroop cost), and Working Memory (3-back d’) as predictors, explained 20% of the variance (R^2^ =0.20). (C) The Adjusted R^2^ of the regressions models to predict EM performance. The variance explained improves with the addition of Stroop Cost to age, but not with the addition of 3-back d’. (D) Predicted versus observed scores for the Energetic Masking (EM) task. The final model included Age and Inhibitory Control only, as the addition of Working Memory did not significantly improve model.

For the informational masking (IM) model, the initial model including age alone was significant. Adding Stroop cost in the second step significantly improved model fit, indicating that inhibitory control contributed unique variance beyond age. Incorporating 3-back d′ in the final step further improved the model, yielding a final R^2^ = 0.20 (Fig. 3A). Predicted IM scores from the final model closely tracked observed IM scores (Fig. 3B).

For the energetic masking (EM) model, adding Stroop cost to the baseline model with age improved predictive performance and increased explained variance. However, the subsequent inclusion of 3-back *d′* did not further enhance the model, as reflected by a decrease in adjusted R^2^ (Fig. 3C). The best-fitting model for EM performance therefore included age and Stroop cost (p< 0.001) as predictors (Fig. 3D).

Overall, these results demonstrate that executive function, particularly inhibitory control, plays a critical role in suprathreshold listening conditions both with informational and energetic masking, while working memory plays a significant role in suprathreshold listening selectively in informational masking alone.

## Discussion

The study investigated how executive function and age contribute to speech perception in noise under two distinct masking conditions: Energetic masking which primarily taxes sensory encoding, and Informational masking which places greater demands on cognitive control. The central finding is that executive functions, specifically working memory and inhibitory control, are powerful predictors of speech-in-noise performance, but their respective influences depend on the nature of the listening challenge. Performance on the cognitively demanding IM task was significantly predicted by both working memory and inhibitory control task, whereas performance on the sensory driven EM task was only significantly predicted by inhibitory control. In both cases, executive functioning measures explained unique variance after controlling for age, suggesting that speech perception in noise draws on distinct cognitive resources in addition to sensory coding fidelity.

### Age-related changes in executive function

We found no association between age and the d’ for the 3-back test of working memory, contrary to prior studies reporting age-related declines on working memory (Bonetti et al., n.d.; Humes et al., 2022). This discrepancy may reflect sampling bias inherent to online data collection – older adults capable of completing an online experiment may represent a more cognitively intact subset of the population. The technological and attentional demands of web-based testing likely selected for individuals with higher baseline cognitive engagement, attenuating observable age effects.

In contrast, inhibitory control as measured by Stroop cost declined significantly with age. This finding is consistent with other studies that identify age-related difficulties in suppressing irrelevant information (Dey & Sommers, 2015; Mutter et al., 2006). The result is also in strong agreement of research linking inhibitory control to speech perception (Dey & Sommers, 2015; Gómez-Álvarez et al., 2023; Knight & Heinrich, 2019). Our results extend this relationship by demonstrating that inhibitory control influences both EM and IM tasks. This suggests that the ability to suppress distraction and maintain attentional focus constitutes a general cognitive resource critical for challenging listening, regardless of the type of masking.

### Differential contributions of working memory and inhibitory control

An important implication of our study is that executive function should not be treated as a ‘monolith’ in the context of speech-in-noise perception. Rather, distinct subcomponents contribute selectivity depending on the distinct demands of the listening task. Working memory was a strong, significant predictor for performance on the IM but not the EM task. The IM task requires the listener to hold a semantically unpredictable sentence in memory, integrating information over time and segregating it from competing talkers, a process highly reliant on working memory (Akeroyd, 2008). In contrast, the EM task involves identifying single, isolated words, which places a minimal load on working memory.

Inhibitory control, in contrast, was a significant predictor of performance on both the EM and IM tasks. This suggests that the ability to suppress distractors and have attentional control is a more important and fundamental resource. Inhibitory control appears to be critical for a broad array of challenging listening environments, regardless of whether interference arises at the sensory or cognitive level.

Although age and executive function measures significantly predicted speech-in-noise performance, the amount of variance explained by the combined regression models was at most 20%. This shows that a vast majority of the performance in speech-in-noise tasks are influenced or driven by factors other than age and these two specific executive functioning measures. Other cognitive factors, or sensory deficits such as cochlear synaptopathy (Bramhall et al., 2025; Garrett et al., 2025; Kujawa & Liberman, 2009; Zink et al., 2024) or impaired fine structure coding (Borjigin & Bharadwaj, 2025; Füllgrabe et al., 2014; Parthasarathy et al., 2020; Zhen et al., 2025), which our study was not designed to measure could play a role in identifying these latent contributors.

### Advantages and considerations of web-based testing

Our study used a large-scale, web-based experiment for its implementation. Auditory psychoacoustic tests have historically been confined to controlled laboratory environments due to concerns over calibration, stimulus delivery, and audiometric screening. Our study, which was built upon the framework of Mok et al 2023, was designed to overcome these challenges. We implemented validated headphone checks and hearing screening procedures which enabled rapid recruitment of a large (N=200) and demographically diverse sample across a wide age range (18-73 years). The success of this design underscores the potential of online paradigms for expanding accessibility, statistical power, and ecological validity in auditory research.

Nonetheless, several limitations merit consideration. Audiometric thresholds were not directly measured but inferred from a speech-in-noise screener. This screener has been validated previously with in-lab comparisons and selects normal audiograms with 90% accuracy. However, since this screener itself involved a speech-in-speech task, it may have additionally preferentially selected for participants with robust temporal fine structure coding and attentional abilities. This could also explain some of the null findings, such as the lack of a significant age-related decline in working memory. However, this potential selection bias also strengthens our positive findings, where we observed significant relationships between executive function and speech-in-noise tasks despite this potentially high-performing, ‘cognitively intact’ cohort.

## Conclusions and clinical implications

In conclusion, our study provides evidence that age-related changes in speech-in-noise perception arise not solely from age-related sensory deficits but as a result of multi-system processes requiring both auditory and suprasensory cognitive resources. Inhibitory control emerges as a general resource that is engaged during many challenging listening conditions, while working memory is a specialized resource required for complex and demanding tasks involving informational masking and longer target sequences. These results underscore the need to expand clinical hearing assessments beyond audiometric and sensory measures to include cognitive evaluations that may additionally capture real-world speech-in-noise difficulties. Finally, our work, conducted on a large and diverse online cohort, demonstrates the feasibility and power of web-based psychoacoustics to tackle complex questions in hearing science.

## Statements and Declarations

### Ethical Considerations

All procedures were approved by the institutional review board of the University of Pittsburgh (STUDY24050210).

### Consent to Participate

All procedures were approved by the institutional review board of the University of Pittsburgh (STUDY24050210).

### Consent for publication

Human subjects provided consent to publish their de-identified data.

### Consent to participate

All subjects provided affirmative informed consent.

### Author contributions

SB – data analysis, stats, writing, CD, EB, GC, CS, AY – Conceptualization, data collection, XZ – statistical analysis, HB – conceptualization, editing, AP – conceptualization, writing

### Data availability

Data and code used in this manuscript are available in Open Science Framework - https://osf.io/x4kgy/

